# Targeted regulation of episomal plasmid DNA expression in eukaryotic cells with a methylated-DNA-binding activator

**DOI:** 10.1101/2021.11.01.466616

**Authors:** Isioma Enwerem-Lackland, Eric Warga, Margaret Dugoni, Jacob Elmer, Karmella A. Haynes

## Abstract

**Purpose:** Targeted regulation of transfected extra-chromosomal plasmid DNA typically requires the integration of 9 - 20 bp docking sites into the plasmid. Here, we report an elegant approach, The Dpn Adaptor Linked Effector (DAL-E) system, to target fusion proteins to 6-methyladenosine in GATC, which appears frequently in popular eukaryotic expression vectors and is absent from endogenous genomic DNA. Methods: The DNA-binding region from the DpnI endonuclease binds 6-methyladenosine within the GATC motif. We used a Dpn-transcriptional activator (DPN7-TA) fusion to induce gene expression from transiently transfected pDNAs.

**Results:** We validated methylation-dependent activity of DPN7-TA with a panel of target pDNAs. We observed stronger transactivation when GATC targets were located upstream of the transcriptional start site in the target pDNA. Conclusion: DAL-E, consisting of a 108 aa, 12 kD DNA-binding adaptor and a 4 bp recognition site, offers a genetically-tractable, tunable system that can potentially be redesigned to recruit a variety of regulators (e.g. activators, silencers, epigenome editors) to transfected plasmid DNA.

**LAY SUMMARY:** Transfection of plasmid DNA (pDNA) is a commonly used method for introducing exogenous genetic material into mammalian cells. Once introduced into cells not all pDNAs express this genetic material at sufficient levels. Current techniques to improve transgene expression are limited and are not always feasible for all plasmids. This report presents a new method to improve gene expression from pDNA. The Dpn Adaptor Linked Effector (DAL-E) binds to methylated adenines in the pDNA resulting in increased expression. This technique has exciting implications for improved genetic engineering of mammalian cells.

**GRAPHICAL ABSTRACT:** 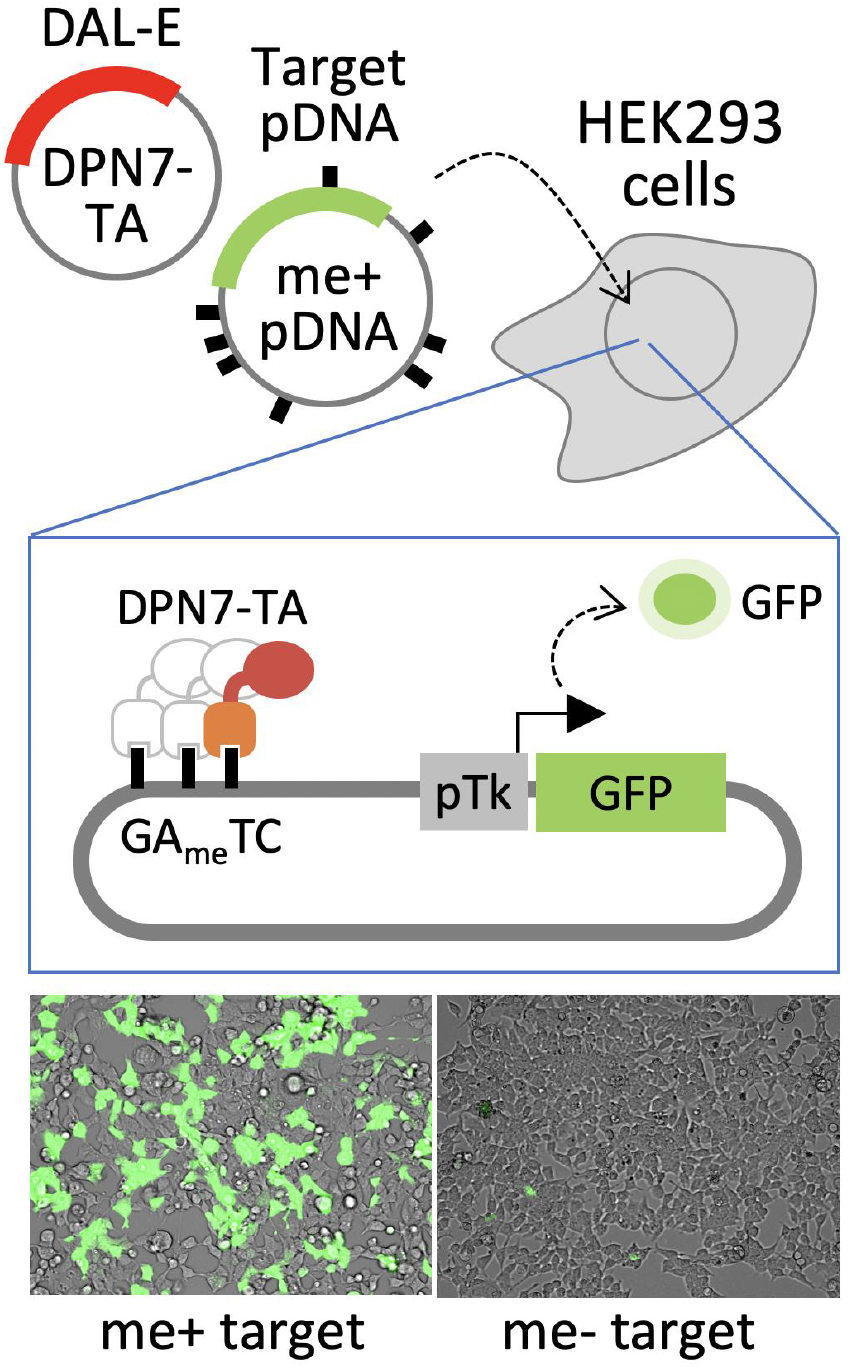

## INTRODUCTION

For many cell research and engineering applications, non-viral plasmid DNA (pDNA) serves as a genetic template that relies upon the cell’s transcriptional machinery to generate molecular products (e.g. proteins and functional RNAs). Although transfection of pDNA with polyplex and lipoplex carriers is a routine practice, optimization is often hampered by inefficient or undetectable expression. A combination of factors such as pDNA size and sequence composition, chemical properties of the delivery vehicle,[1–3] and endogenous transcriptional silencing [4] can impact transgene expression. Furthermore, suboptimal expression can be compounded by co-expressing two or more pDNAs, which is often required for therapeutic adeno-associated virus (AAV) particle production.[5]

To gain control of pDNA expression, transcription-regulating features can be added to the pDNA. For instance, we have demonstrated that adding NF-Y and MYB transcriptional activator target DNA sequences upstream of a minimal promoter increased luciferase expression from transfected pDNA in prostate cancer PC-3 cells.[6,7] Fusion proteins can also be used as transgene-specific regulators to boost expression in mammalian cells.[8–11] Both approaches, enhancer elements and fusion protein regulators, require the addition of DNA target sequences into the pDNA, which may not always be possible or desirable.

Here, we present a new tool called “Dpn adaptor linked effector” or DAL-E (named for Marie M. Daly, biochemist) to target fusion proteins, such as regulators, to transfected pDNA. DAL-E targets the DNA methylation signal 6-methyladenosine (6-meA) at the GATC motif (GA_me_TC) [12], which is present in bacteria but absent from eukaryotic DNA. This four base pair motif has a smaller footprint than other regulator binding sites (e.g., 9 bp for zinc fingers), and can be added or eliminated from pDNA via site directed mutagenesis. The restriction enzyme DpnI exclusively binds GA_me_TC. Barnes et al. used DpnI, with Mg^2+^ to inhibit nuclease activity, to isolate bacterial DNA for metagenomic analysis from saliva samples that contained prokaryotic and mammalian DNA.[13] Later, Kind et al. eliminated the nuclease activity of DpnI by truncating it into a shorter protein, DPN7 (109 amino acids), which enabled localization of a fused green fluorescent protein to artificially generated GA_me_TC sites within mammalian chromosomal DNA.[14] A DPN7-VP64 fusion has been used to model chromatin-templated epigenetic mechanics in HEK-293 chromosomal DNA.[15] In our current study, we investigated whether DPN7-VP64 could regulate genes on extrachromosomal pDNA, and how the number and positioning of GA_me_TC along the plasmid affects the activity of DPN7-VP64.

## RESULTS

### A DPN7-VP64 fusion specifically mediates *trans*-activation of 6-methyl-adenosine enriched plasmid DNA in mammalian cells

Typically, artificial transactivation of genes is controlled by transcriptional activation domains fused to DNA-binding domains such as Gal4, zinc fingers (ZFs), and dCas9/gRNA complexes that are designed to bind to a specific sequence at a precise, optimal location relative to the core promoter. In contrast, DPN7 binding sites (GATC sequences) can appear at multiple sites across the gene or plasmid. To determine if the broad binding of DPN7 would support or hinder transactivation, we compared the ability of a DPN7-transactivator fusion (DPN7-TA) and a Gal4 DNA-binding transactivator fusion (Gal4-TA) (Supplemental Fig. 1A) to enhance the expression of a fluorescence gene from methylated plasmid DNA. The target plasmid UAS-pTk-CFP contains tandem copies of a core Gal4 upstream activation site (UAS) sequence (5x 5’-cggagtactgtcctccga) upstream of a herpes simplex virus thymidine kinase core promoter (HSV pTk) (Supplemental Fig. 1A). UAS-pTk-CFP has 23 GATC sites; three of these GATC sites are located within 500 bp of the pTk transcription start site (TSS) at -452, -460, and -468 (Supplemental Fig. 1B). To generate methylation-free or methylation-enriched plasmids, we propagated the target plasmid in either a dam-/dcm- or dam+/dcm+ strain (respectively) (Supplemental Fig. 1C).

We cotransfected HEK-293 cells with plasmids that express a transactivator and either a methylated or unmethylated CFP target reporter (Fig. 1A,B). In a control experiment designed to confirm that VP64 could stimulate transcriptional activation when bound near pTk, we observed similar levels of Gal4-TA-mediated activation of unmethylated (me-) and methylated (me+) UAS-pTk-CFP as shown by the percent of CFP positive cells (0.88-fold me+/me-, p=0.66) and mean CFP signal intensity (0.71-fold me+/me-, p=0.12) (Fig. 1C, D).

**Figure 1.**
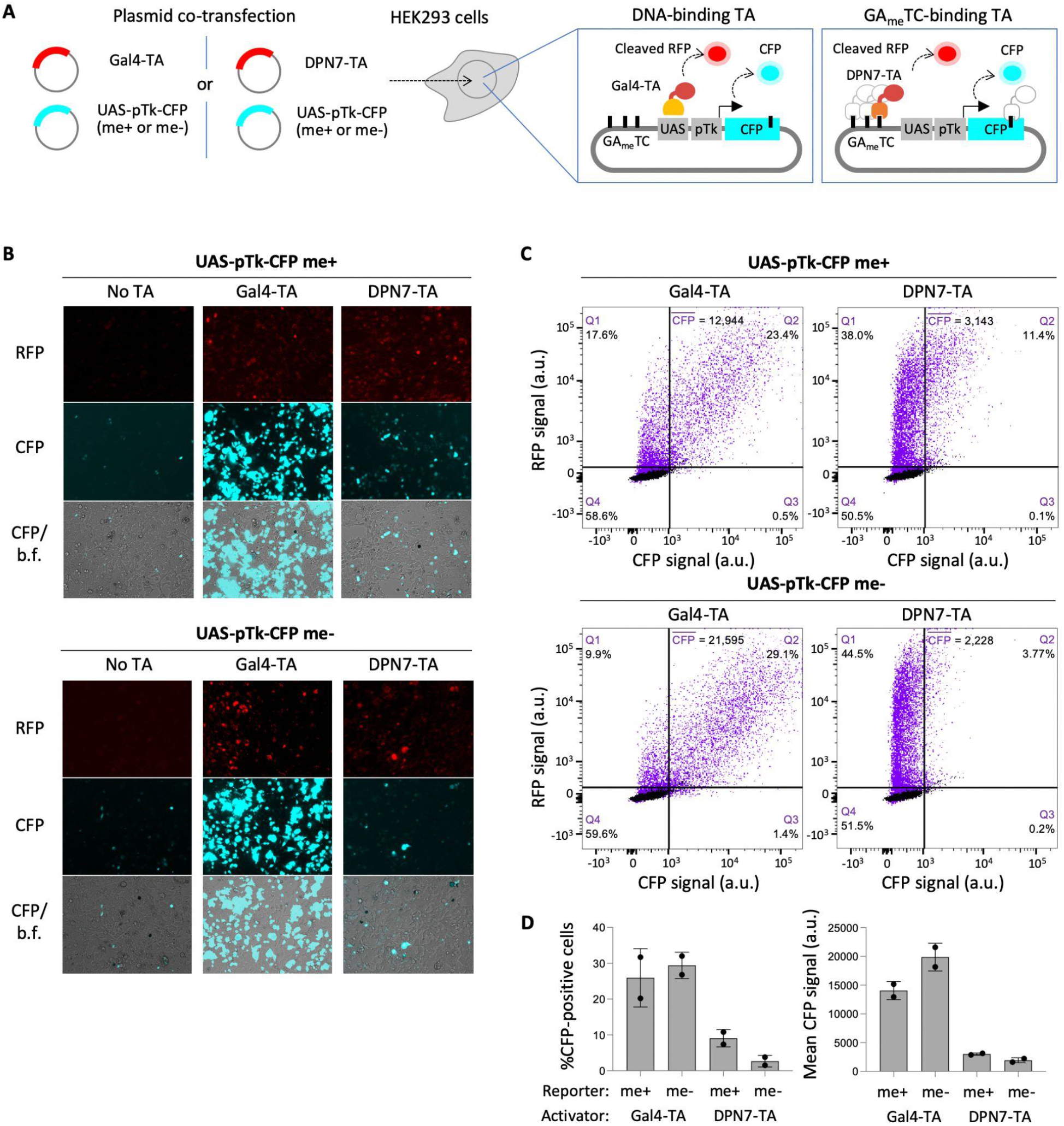
DPN7 interacts with GA_me_TC-enriched, transfected pDNA. (A) Experimental design to compare regulation by a single target site binding transactivator (Gal4-TA) versus the GA_me_TC-binding transactivator (DPN7-TA) that potentially binds at pTk-proximal (solid filled) and distal sites (grey outlined). (B) Representative fluorescence microscopy images of transfected HEK293 cells. (C) A total of 20,000 cells were analyzed per construct. Scatter plots of flow cytometry data from live gated co-transfected cells described in panel A. Black = untransfected cells, purple = co-transfected cells. (D) Bar graphs show average values of two replicate transfections: %CFP-positive cells (left) and arithmetic mean signal value of the CFP-positive populations (right). Error bars indicate standard deviation. Unpaired t-test used to determine significance, p values reported in text.

When DPN7-TA was co-expressed with the methylated and unmethylated target reporter we observed a small increase in both percent of CFP positive cells (3.39-fold me+/me-, p=0.1) and mean CFP signal (1.57-fold me+/me-, p=0.13), with the methylated reporter (Fig. 1C,D). These results suggest that DPN7-TA activates reporter expression in a methylation-dependent manner. GAL4-TA showed the highest overall activation of the CFP reporter (No TA vs. GAL4-TA) (Fig. 1 B,C). These results prompted us to consider the differences in the engagement of the transactivator proteins with the target pDNA. Gal4-TA binds to the UAS sites upstream of the promoter, while the DPN7-TA could potentially bind to 23 different GA_me_TC sites positioned throughout the plasmid. We thus hypothesized that the position and number of GA_me_TC could have an effect on the robustness of DPN7-TA mediated activation.

### *Trans*-activation via DPN7-TA is strongest for plasmids with TSS-proximal GATC sites

We sought to determine if the number and positioning of GATC sites relative to the transcription start site affects DPN7-TA activity. We co-transfected the DPN7-TA plasmid with GFP-expressing target plasmids that carried 9 to 21 GATC sites at different positions (Fig. 2A). For these experiments, we used a smaller (∼4000 bp) vector that was amenable to the removal and addition of GATC sites, and carried the same promoter as the UAS-pTK-CFP plasmid from our previous experiments. pTk-GFP-21 has 21 GATC sites; four of these GATC sites are located within 500 bp of the pTk transcription start site (TSS) (Fig. 2B). Comparing the effect of the DPN7-TA on the methylated and unmethylated pTk-GFP-21 reporter sites, we measured a modest increase in the percent of GFP positive cells (1.8-fold me+/me-) and mean GFP signal intensity (2.1-fold me+/me-), further demonstrating that DPN7-TA recognizes methylated GATC sites on the target plasmid (Fig. 2C).

**Figure 2.**
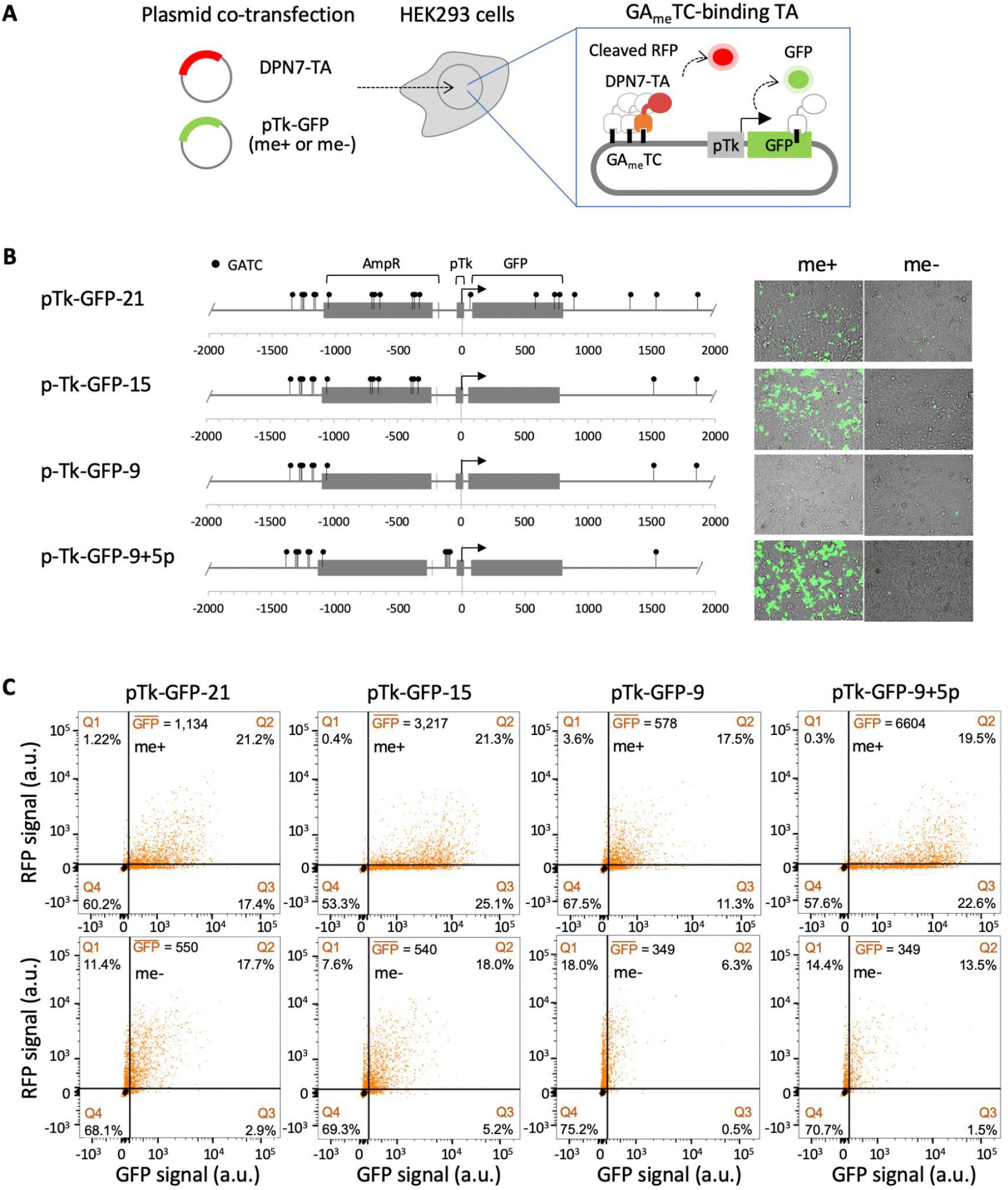
Position of GATC sites relative to the TSS affect DPN7 mediated pDNA activation (A)Experimental design to observe the effects of varying GATC site number and position on DPN7-TA mediated pDNA expression. (B) The plasmid maps show GATC positions relative to the transcription start site for each target plasmid used in this section of our study. (C) A total of 20,000 cells were analyzed per construct. Scatter plots of flow cytometry data from live gated co-transfected cells described in panel A (data represents a single replicate experiment). Black = untransfected cells, orange = cotransfected cells.

Next, we investigated whether the removal of GATC sites from the open reading frame (ORF) of the reporter would enable stronger DPN7-TA-mediated GFP expression. We tested plasmid pTk-GFP-15, where 6 GATC sites were removed downstream of the TSS, 3 from the GFP ORF (Fig. 2B). Again, DPN7-TA co-expression increased GFP expression from the methylated pTk-GFP-15 reporter, compared to the unmethylated reporter (2.83-fold me+/me-). This fold increase, in GFP mean signal intensity, was higher than what we observed with the pTk-GFP-21 methylated reporter (Fig. 2C). These results suggest that GATC sites within 1000 bp downstream of the TSS or sites within the transcribed ORF could have inhibitory effects on the activation of plasmid expression by DPN7-TA.

To assess the importance of methylation sites 1000 bp upstream of the TSS, 6 of the 7 GATC sites in the ampicillin resistance gene (AmpR) were removed, resulting in a pTk-GFP-9 reporter with a total of 9 GATC sites (Fig. 2B). We observed a reduction of mean signal intensity with the methylated pTk-GFP-9 reporter (578 a.u.) when compared to the pTk-GFP-21 (3217 a.u.) and pTk-GFP-15 (1134 a.u.) reporters. This reduction of GFP signal intensity suggests that the presence of methylated GATC sites 1000 bp upstream of the TSS is important for DPN7-TA-mediated transcriptional activation of pTk-GFP.

To further test this idea, we added 5 GATC sites 24 bp upstream of the promoter in pTk-GFP-9. With the resulting reporter, pTk-GFP-9+5p, we observed an increase in GFP positive cells when the DPN7 transactivator was co-expressed with the methylated reporter. Most interestingly, when comparing the methylated pTk-GFP-9+5p reporter to every other methylated reporter, we observed the most intense GFP signal as evidenced by our cell images and from the mean signal intensity data (6604 a.u.) (Fig. 2B,C). This allowed us to conclude that the positioning of methylation sites relative to the TSS is extremely important for DPN7-TA mediated plasmid regulation. It is clear that a removal of proximal GATC sites downstream of the TSS and the addition of proximal GATC sites upstream of the TSS has the most desirable effects on the transgene expression when DPN7-TA is coexpressed in the host cells. Future studies looking at varying the number of sites in the 5xGATC cluster, as well as removing all other sites besides this cluster would potentially result in an even more fine tuned system.

## DISCUSSION AND CONCLUSION

We conclude that DPN7-TA activates expression from pDNA in a methylation-dependent manner, and that the positioning of methylation sites relative to the TSS affects DPN7-TA activity. We have shown that removal of proximal GATC sites downstream of the TSS and addition of proximal GATC sites upstream of the TSS can increase DPN7-TA-driven transcriptional activation. Our results and work from other groups [14,15] provide strong evidence that the GA_me_TC motif from prokaryotic DNA can be selectively targeted within live eukaryotic cells by a DpnI-derived module.

Plasmid DNA transfection is a cornerstone method for cell biology research and cell engineering, but efficient expression of synthetic genes remains a barrier. DAL-E provides a new method for transactivation to overcome dampened transgene expression. The DNA binding footprint and the DPN7 module (324 bp, 108 aa, 12 kD) are small and amenable to DNA construction and expression. Furthermore, GATC sites often appear naturally in popular plasmid vectors. While the DPN7-VP64 used in this study activates gene expression by recruiting endogenous transcription factors from the transcription initiation complex [16], transactivation could be modulated with chromatin modifying enzymes. A DPN7-histone acetyltransferase (e.g. p300) fusion could be used to modify under-acetylated histones that dampen pDNA expression [17]. DAL-E can potentially be modified to suppress expression by coupling DPN7 with a repressor (e.g. KRAB). DAL-E presents new opportunities to control gene expression from pDNA to support gene delivery and cell engineering.

## MATERIALS AND METHODS

### Construction and methylation of recombinant DNA

Sub-parts (modules, Supplemental Table 1) were amplified via high-fidelity polymerase chain reaction (PCR) (New England Biolabs #M0491S) using dsDNA templates and ssDNA primers listed in Supplemental Table 2. To construct DPN7-VP64-RFP (KAH252_MV10) and Gal4-VP64-RFP (KAH253_MV10), we used “BioBrick” digestion/ ligation assembly [18] standard RFC23 [19] (see Supplemental Tables 1 and 2). KAH252 and KAH253 were each digested with XbaI/ SpeI and cloned into the XbaI site of MV10, between 5’ CMV promoter and Kozak, and a 3’ NLS-6xhis-stop sequence [20]. The CFP reporter UAS-pTk-CFP (BD001_MV2) was constructed using BioBrick Assembly. The GFP reporter pTk-GFP-21 was constructed from pEF-GFP (Addgene #11154). The EF1a promoter was replaced by a SalI/ KpnI pTk promoter fragment. GATC sites were sequentially removed by cloning in synthetic DNA fragments with mutated GATC sites. To construct pTk-GFP-9+5p, a 5xGATC annealed oligo fragment was inserted into pTk-GFP-9 at a XbaI site 24 bp upstream of the pTk promoter.

The CFP and GFP reporter plasmids were propagated in either dam-/dcm-strain C2925 (New England Biolabs) or dam+/dcm+ strain DH5-alpha Turbo (NEB C2984) to produce unmethylated or methylated plasmids, respectively. Methylation was analyzed by DNA gel electrophoresis of 600 ng of DpnI- or MboI-digested plasmid DNA and GeneRuler 1 kb DNA Ladder (Thermo Fisher SM0313). All annotated plasmid sequences for the transactivator- and reporter-expressing plasmids are available in the Haynes Lab Benchling database (https://benchling.com/hayneslab/f_/mrI2t4mM-collection-dal-e/).

### Cell culture and transfection

HEK293 cells were cultured in Dulbecco’s Modified Eagle Medium supplemented with 10% fetal bovine serum and 1% penicillin and streptomycin. Cells were grown at 37°C in a humidified CO_2_ incubator. Transfection of fluorescent reporters and transactivators were done in 6-well plates. 3×10^5^ cells were seeded, grown to 80% confluency, and transfected with plasmid DNA/ lipofectamine complexes (Lipofectamine LTX (Life Technologies, Carlsbad, CA), 2.5 ug total DNA) per manufacturer’s instructions. Co-transfections included 250 ng Gal4-TA or DPN7-TA plasmid and 2.25 ug CFP or GFP reporter plasmid. No transactivator (No TA) transfections included 2.5 ug of CFP or GFP reporter plasmid. Cells were imaged and harvested for flow cytometry 72 h post transfection.

### Imaging

Cells were imaged using EVOS M5000 (Invitrogen, Waltham, MA). Image overlays were generated from tiff files converted to 8 bit grayscale, and processed with the Merge Channels tool in the FIJI/ ImageJ version 1.53k software package.

### Flow cytometry

Transfected cells were harvested using Accutase cell dissociation solution (Sigma Aldrich, St Louis, MO) and counted with TC20 Automated cell counter (Biorad, Hercules, CA). Approximately 1×10^6^ cells were resuspended in 500 µl of sorting buffer (1xPBS (Ca/Mg++ free), 5mM EDTA, 1%BSA, 25mM HEPES, filter sterilized). Cells were then sorted on a BD FACSymphony A5 flow cytometer at the Emory Flow Cytometry Core (EFCC). Flow cytometry data was analyzed using FlowJo software version 10.

## DECLARATIONS

This project was supported by the Emory/ Georgia Tech Wallace H. Coulter Department of Biomedical Engineering, and in part by the Emory Flow Cytometry Core (EFCC, an Emory Integrated Core Facility, EICF) and the National Center for Georgia Clinical & Translational Science Alliance of the National Institutes of Health (Award UL1TR002378 to Emory University). The content is solely the responsibility of the authors and does not necessarily reflect the official views of the National Institutes of Health. The authors declare no competing interests. Recombinant DNA sequences are available online at https://benchling.com/hayneslab/f_/mrI2t4mM-collection-dal-e/.

