## Supplementary Information for "Targeted regulation of episomal plasmid DNA expression in eukaryotic cells with a methylated-DNA-binding activator"

*bioRxiv*

Isioma Enwerem-Lackland[1], Eric Warga[2], Margaret Dugoni[2], Jacob Elmer[2], and Karmella A. Haynes[1]

1. Wallace H. Coulter Department of Biomedical Engineering, Emory University, Atlanta, GA 30322
2. Department of Chemical & Biological Engineering, Villanova University, Villanova, PA 19085

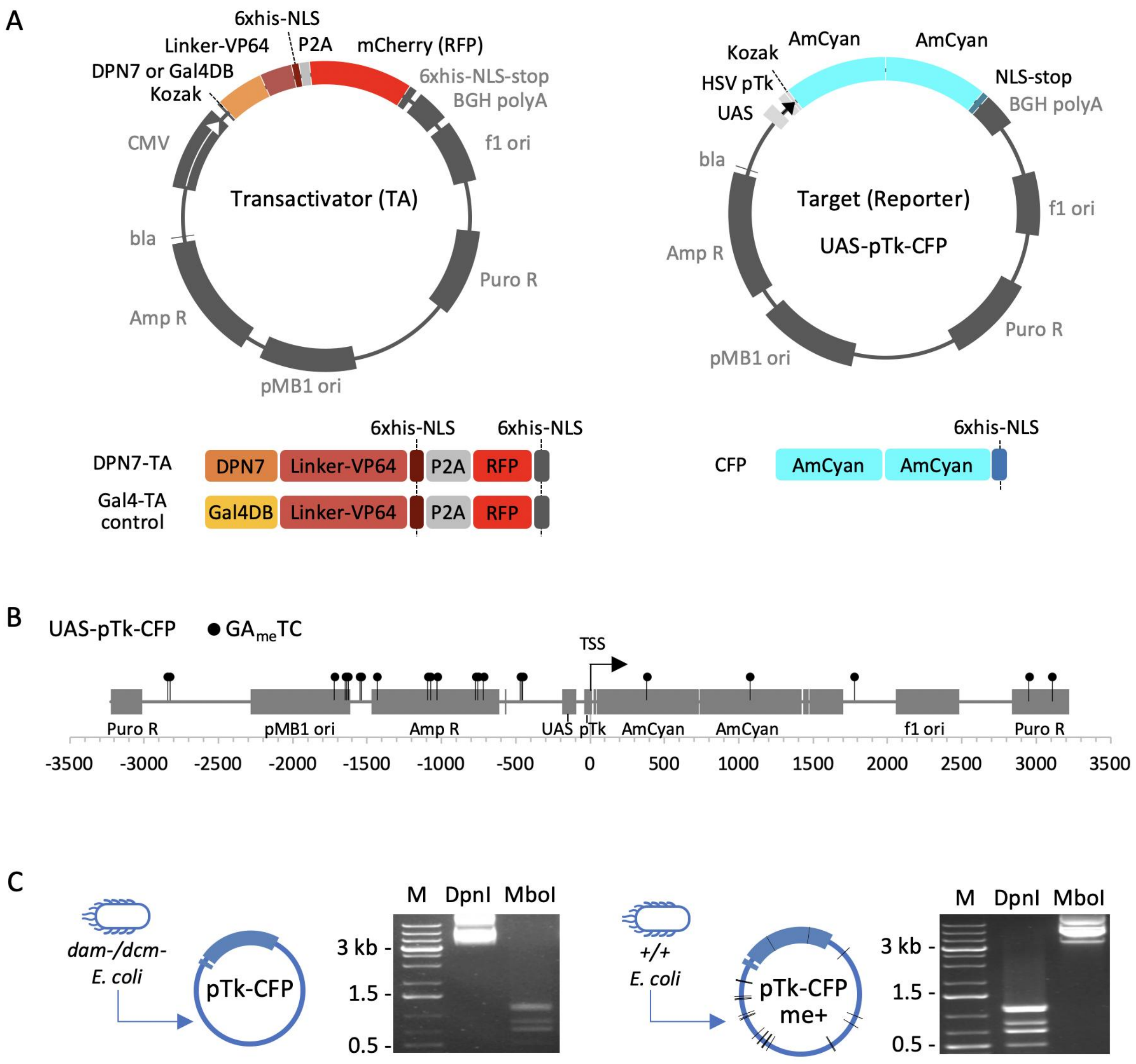


**SUPPLEMENTAL FIGURE 1.** Constructs and target reporter methylation. (A) Plasmid DNA maps (top) and protein domain maps (bottom). The transactivators include a VP64 transactivation domain and either an N-terminal DPN7 domain or Gal4 DNA binding domain (Gal4DB). P2A-mCherry was included to co-express a cleaved red fluorescent protein (RFP). The target reporter plasmid UAS-pTk-CFP expresses a 2x fusion AmCyan protein. NLS = nuclear localization signal. (B) Map of the 23 GATC sites in plasmid UAS-pTk-CFP. (C) Digests with DpnI (6-meA-dependent) and MboI (6-meA-blocked) confirmed methylation states of control and experimental target plasmids. M = "marker" 1 kb DNA ladder.

**SUPPLEMENTAL TABLE 1. Sequences of DNA fragments used to build the DPN7-TA and Gal4-TA regulators.**

| **Module** | **DNA Sequence (5’ … 3’)** |
|---|---|
| DPN7 | ccttctaaaggaaggatatttcttgtgcaagatggacaagttagagatccagaaaaagttacaaaagaatttaagcaaggtttattttt aaggaagagctctctgtcatcaagaggttggacaatagaaattctaaattgtatagataagatagagggttcagaatttacccttga agatatgtatcgttttgaaagtgacctaaaaaatatctttgttaagaacaatcatatcaaagaaaagattaggcaacagcttcaaata ttaagagacaaagaaataatagaatttaaaggtagaggaaagtatcggaaattattcgaa |
| Gal4DB | atgaagctactgtcttctatcgaacaagcatgcgatatttgccgacttaaaaagctcaagtgctccaaagaaaaaccgaagtgcgc caagtgtctgaagaacaactgggagtgtcgctactctcccaaaaccaaaaggtctccgctgactagggcacatctgacagaagt ggaatcaaggctagaaagactggaacagctatttctactgatttttcctcgagaagaccttgacatgattttgaaaatggattctttaca ggatataaaagcattgttaacaggattatttgtacaagataatgtgaataaagatgccgtcacagatagattggcttcagtggagact gatatgcctctaacattgagacagcatagaataagtgcgacatcatcatcggaagagagtagtaacaaaggtcaaagacagttg actgtatcgccggaatttccggggatc |
| Peptide linker (GS) | ggcggtggcggatctggaggtggtggttcaggcggaggcggatctggaggaggtggttca |
| VP64 Activation domain | gacgctttggacgacttcgacttggacatgttgggttctgacgctttggacgacttcgacttggacatgttgggttctgacgctttggacg acttcgacttggacatgttgggttctgacgctttggacgacttcgacttggacatgttg |
| Nuclear localization signal | cccaagaaaaagcgcaaggta |
| 6xhis epitope and P2A cleavage | caccatcaccaccatcacactagaggcagtggagctactaacttcagcctgctgaagcaggctggtgacgtcgaggagaatcct ggcccc |
| mCherry (RFP) | gtgagcaagggcgaggaggataacatggccatcatcaaggagttcatgcgcttcaaggtgcacatggagggctccgtgaacgg ccacgagttcgagatcgagggcgagggcgagggccgcccctacgagggcacccagaccgccaagctgaaggtgaccaagg gtggcccccctgcccttcgcctgggacatcctgtcccctcagttcatgtacggctccaaggcctacgtgaagcaccccgccgacatc cccgactacttgaagctgtccttccccgagggcttcaagtgggagcgcgtgatgaacttcgaggacggcggcgtggtgaccgtga cccaggactcctccttgcaggacggcgagttcatctacaaggtgaagctgcgcggcaccaacttcccctccgacggccccgtaat gcagaagaagaccatgggctgggaggcctcctccgagcggatgtaccccgaggacggcgccctgaagggcgagatcaagc agaggctgaagctgaaggacggcggccactacgacgctgaggtcaagaccacctacaaggccaagaagcccgtgcagctac ccggcgcctacaacgtcaacatcaagttggacatcacctcccacaacgaggactacaccatcgtggaacagtacgaacgcgcc gagggccgccactccaccggcggcatggacgagctgtacaag |

**SUPPLEMENTAL TABLE 2. Sub-parts used to build expression plasmids for DPN7-TA and Gal4-TA.** For parts that were clones via PCR, primers were synthesized by Integrated DNA Technologies (IDT). Five prime extensions (lowercase) added a 5' XbaI site and 3' SpeI (BcuI), NotI, and PstI sites to each module (sub-part). Unless otherwise noted, PCR-amplified modules were digested with XbaI and PstI, and then ligated into V0120 (AmpR, pMB1) at XbaI and PstI sites. The final plasmids are listed in the final column. Plasmid names and ID's are from the Haynes lab Benchling database online. Please use these names and ID's when requesting plasmid DNA. To construct DPN7-TA (KAH252), step-wise digestion/ligation assembly (BioBrick standard RFC23 [19]) was used to build the following, in order: NLS-6xhis-P2A (KAH241), DPN7-PLflex (KAH242), DPN7-PLflex-VP64 (KAH248), DPN7-PLflex-VP64-NLS-6xhis-P2A (KAH250), then DPN7-PLflex-VP64-NLS-6xhis-P2A-mCh (KAH252). For Gal4-TA (KAH252) we built the following, in order: Gal4DB-PLflex (KAH247), Gal4DB-PLflex-VP64 (KAH249), Gal4DB-PLflex-VP64-NLS-6xhis-P2A (KAH251), then Gal4DB-PLflex-VP64-NLS-6xhis-P2A-mCh (KAH253).

| Module | PCR Template | Forward Primer (5'-) | Reverse Primer (5'-) | Final plasmid |
|---|---|---|---|---|
| DPN7: GAmeTC binding | pHIV-Tet-Puro-IRES-EGFP-DPN7 [14] | cctttctagaccttctaaaggaaggatatttcttgt | aaggctgcagcggccgctactagtttcgaataatttccgatactttcct | DPN7_V0120 (KAH025.2) |
| Gal4DB: UAS binding | Prior subclones | cctttctagaatgaagctactgtcttctat | aaggctgcagcggccgctactagtgatccccggaaattccggcg | Gal4DB_V0120 (DT02) |
| PLflex: peptide linker | n/a (synthesized DNA) | n/a | n/a | PLflex_V0120 (KAH021) |
| VP64: transcriptional activation | n/a | n/a | n/a | VP64_V0120 (C0070) |
| NLS: Nuclear localization tag | n/a (annealed ssDNA oligos) | n/a | n/a | NLS_V0120 (T0100) |
| 6xhis-P2A: epitope and post-translation cleavage | n/a (annealed ssDNA oligos) | n/a | n/a | 6xhis-P2A_V0120 (DBN030) |
| mCh: mCherry RFP | n/a (prior subclone) | n/a | n/a | E3000_V0120 |
